# Advancements in Ligand-Based Virtual Screening through the Synergistic Integration of Graph Neural Networks and Expert-Crafted Descriptors

**DOI:** 10.1101/2023.04.17.537185

**Authors:** Yunchao (Lance) Liu, Rocco Moretti, Yu Wang, Ha Dong, Bailu Yan, Bobby Bodenheimer, Tyler Derr, Jens Meiler

## Abstract

The fusion of traditional chemical descriptors with Graph Neural Networks (GNNs) offers a compelling strategy for enhancing ligand-based virtual screening methodologies. A comprehensive evaluation revealed that the benefits derived from this integrative strategy vary significantly among different GNNs. Specifically, while GCN and SchNet demonstrate pronounced improvements by incorporating descriptors, SphereNet exhibits only marginal enhancement. Intriguingly, despite SphereNet’s modest gain, all three models-GCN, SchNet, and SphereNet-achieve comparable performance levels when leveraging this combination strategy. This observation underscores a pivotal insight: sophisticated GNN architectures may be substituted with simpler counterparts without sacrificing efficacy, provided that they are augmented with descriptors. Furthermore, our analysis reveals a set of expert-crafted descriptors’ robustness in scaffold-split scenarios, frequently outperforming the combined GNN-descriptor models. Given the critical importance of scaffold splitting in accurately mimicking real-world drug discovery contexts, this finding accentuates an imperative for GNN researchers to innovate models that can adeptly navigate and predict within such frameworks. Our work not only validates the potential of integrating descriptors with GNNs in advancing ligand-based virtual screening but also illuminates pathways for future enhancements in model development and application. Our implementation can be found at https://github.com/meilerlab/gnn-descriptor.

## 1. Introduction

Virtual screening is a major way to supplement traditional high-throughput screening (HTS) for cost and time efficient drug discovery^1^. Two major branches of virtual screening exist: ligand-based, and structure-based. For the application of structure-based methods, detailed knowledge of the target’s structure is essential, typically acquired through experimental methods such as X-ray crystallography or nuclear magnetic resonance (NMR). In cases where experimental data is lacking, computational predictions like homology modeling are employed to infer the three-dimensional configurations of targets. Recently, there are many AI-driven protein structure prediction tools available as well, such as AlphaFold ^2^, RosettaFold^3, 4^, ESMFold^5^.

This work focuses on the ligand-based method, for situations where the target structure remains unknown or cannot be computationally predicted. These methods depend on the knowledge of previously identified active compounds that bind to the target, leveraging this information to identify new potential drugs^6^. Even in the age that computational protein structure prediction tools are available, ligand-based approaches are needed for several reasons. First, while structure prediction tools have made remarkable progress, there are still limitations in their ability to accurately predict all protein structures, especially for proteins with highly dynamic regions and transient conformations. The ligand-based method does not require structural information, making it valuable for targets where high-quality structures are not available. Secondly, ligand-based methods can sometimes be faster and less resource-intensive than structure-based methods, especially in the early stages of drug discovery. They allow researchers to quickly screen vast chemical spaces or compound libraries to identify potential hits without detailed structural information. Thirdly, some targets have multiple or flexible binding sites that can be challenging to characterize with structure-based methods alone. Ligand-based methods can help identify ligands that interact with such targets by leveraging data from known active compounds without relying on a fixed 3D structure.

Meanwhile, numerous studies applied GNNs to molecule-related tasks, given the intrinsic graph nature of molecules ^7-13^. While some of those tasks achieve good results, several factors still make GNN for molecule representation learning challenging. First, data available for training in drug discovery campaigns is usually limited due to the high cost of experimental assays. Secondly, GNNs typically have difficulty learning molecular-level features due to their limited receptive field or learning non-additive molecular-level features such as total polar surface area. Thirdly, GNN intrinsically suffers from problems such as over-smoothing ^14^ and over-squashing ^15^ that introduce information loss in obtaining the global learned embedding from the atomic features.

As a solution, integrating the expert knowledge in the GNN workflow has become a new trend ^16^. Expert knowledge can help supplement the data-hungry GNNs with prior knowledge to increase data efficiency and overcome intrinsic GNN shortcomings. One of the simplest ways to integrate expert knowledge is to combine the expert-crafted descriptors with GNN-learned representation through concatenation ^17, 18^. However, while commonly used, a thorough evaluation of this concatenation strategy is lacking.

This work contributes to the field by comprehensively evaluating this commonly used strategy in a virtual screening setting using nine well-curated HTS datasets. We find that although this strategy is often effective, it is not always the case. Additionally, we discover that the combined GNNs show convergence of performance metrics, suggesting the potential interchangeability of sophisticated GNN architectures with simpler counterparts under this integrative strategy. Moreover, surprisingly we found that descriptors are fairly robust under the scaffold split scenario, which is often a more realistic setting in a drug discovery campaign. These findings prompt the need to examine the current integration strategies to understand their limitations, find better ways to integrate domain expert knowledge and provide a path for more advanced ligand-based virtual screening.

## 2. Results

### 2.1. Concatenate descriptors with GNNs

As shown in Figure 1, the concatenation strategy ^17, 18^ examined in this work proposes to train a neural network to predict activity by combining a GNN-derived molecular representation with the expert-crafted descriptors ^19^. Specifically, for a representation *h* from the GNN, it is concatenated with the descriptor *h*_*dp*_.

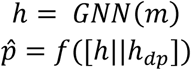

where *m* is the input molecular graph and, *h*_*dp*_ is a descriptor. *f* (·) is a classifier, usually a Multi-Layer-Perceptron (MLP). 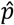 is the predicted activity.

**Figure 1.**
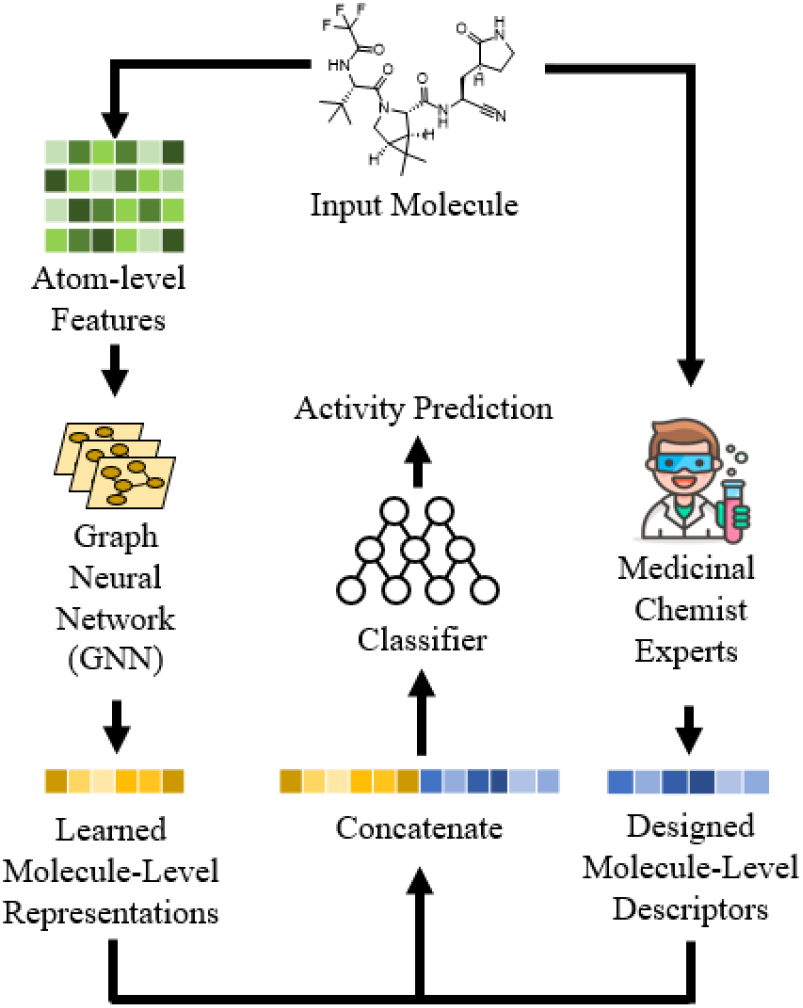
Overview of the investigated method. The learned molecular representation of GNN is concatenated with expert-crafted descriptors to enhance the predictive power.

The model is trained by optimizing the binary cross entropy loss *L*:

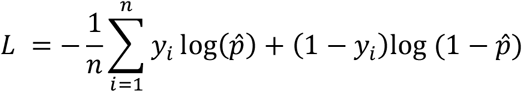

In this work, we used three GNN models in our experiments: GCN ^20^, SchNet ^11^ and SphereNet ^13^. We used the BioChemical Library (BCL) ^21^ to generate descriptors.

where *n* is the number of samples in a batch, and y_i_ is the experimentally determined active/inactive status of the *i*-th molecule.

### 2.2. Effectiveness of the Concatenation Strategy Varies for GNNs with Random Split

In Figure 2 the boxplots of model performances evaluated using four different metrics are shown (Experiments are detailed in Section 3 Method). The p-value is calculated using paired t-test ^22^.

**Figure 2.**
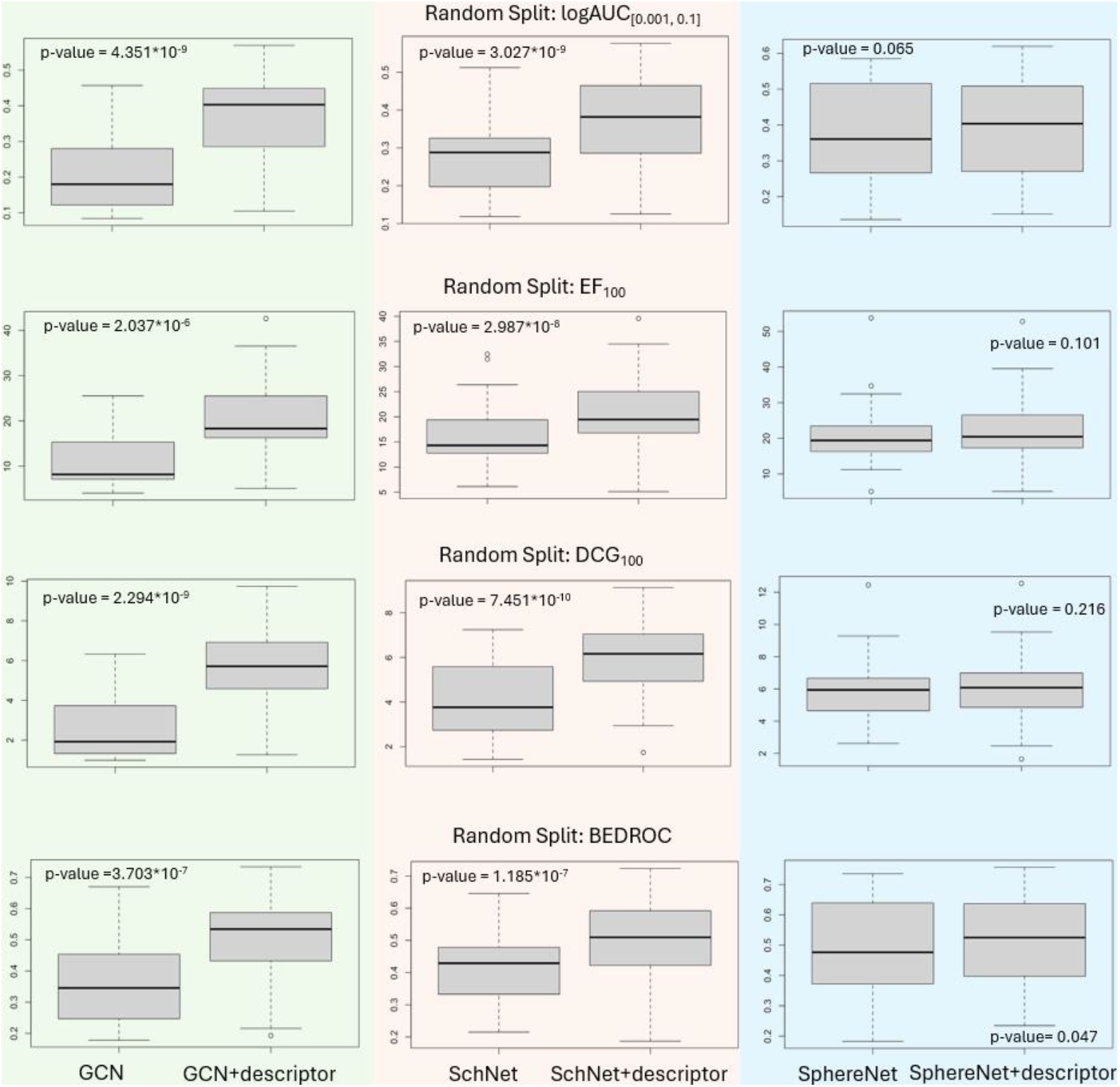
Random Split: Performance of three GNNs with their corresponding descriptor-integrated counterpart.

The significant improvements observed in both GCN and SchNet models across four evaluation metrics highlight the investigated strategy’s potential to facilitate the identification of bioactive compounds in drug discovery. Although the benefits were less pronounced for the SphereNet model (as a bigger p-value is observed), the overall results advocate for the integration strategy’s adoption as a valuable tool in computational chemistry.

There are three rationales for this approach. First, data available for training in drug discovery campaigns is usually limited due to the high cost of experimental assays. The expert-crafted descriptors supplement GNNs with prior knowledge, i.e., descriptors that worked well in virtual screening in the past, which reduces the need for GNNs to learn that knowledge from a large amount of data. Secondly, GNNs typically have difficulty learning molecular-level features due to their limited receptive field or learning non-additive molecular-level features such as total polar surface area. On the other hand, molecular-level descriptors provide global features directly. Thirdly, GNN intrinsically suffers from problems such as over-smoothing ^14^ and overs-quashing ^15^ that introduce information loss in obtaining the global learned embedding from the atomic features. Meanwhile, the descriptors extract the molecular features directly and circumvent information loss, complementing GNN-learned embeddings.

### 2.3. All Descriptor-integrated GNNs Converge to Similar Performance with Random Split

The analysis undertaken in this study revealed significant insight regarding the investigated strategy’s performance. Initially, the GNNs—each with its intrinsic computational complexities and capabilities—demonstrated disparate levels of efficacy. However, upon the integration of descriptors, a notable convergence in their performance metrics was observed, spanning all four evaluated metrics. As shown in Figure 2, SphereNet and SchNet, are more advanced GNNs compared with GCN. Yet, when these advanced GNNs were coupled with descriptors, the resultant performance was not just enhanced but aligned closely with that of their simpler counterparts GCN.

This intriguing outcome underscores the potency of the integration strategy in equalizing the performance landscape among GNN architectures. By integrating expert-crafted descriptors through the integration approach, even less complex GNN models could elevate their predictive accuracies to levels akin to those of more complex GNNs. Essentially, the integration strategy acts as a performance catalyst, diminishing the gaps between GNN models of varying complexities and facilitating a more uniform field of competition. Such findings highlight the potential of combining deep learning techniques with established domain knowledge, suggesting a reevaluation of the necessity for complex GNNs in scenarios where their simpler counterparts can achieve comparable outcomes through integration with descriptors.

### 2.4. Expert-crafted Descriptor Still Outperforms Most GNNs Using Scaffold Split

Besides random split, we also conducted experiments on scaffold split. This is a realistic scenario because medicinal chemists often need to determine the activity of structures substantially different from those in the known training set. They seek these structural differences for various reasons, such as avoiding patented structures, finding simpler synthetic routes, improving compound properties etc ^23^.

As expected, the overall performance under the scaffold split decreased compared with that under the random split. This decrease is due to the greater difficulty in predicting the performance of structures significantly different from the training set, as the data distribution differs between training and testing. However, as shown in Figure 3, the results from the scaffold split evaluation solidify the potential of the integration strategy in enhancing the performance of various GNN architectures for ligand-based virtual screening. The combined GNN-derived molecular representations with descriptors, improve the identification and prioritization of active compounds (Although outliers exist, which is consistent with our results for random split that the effectiveness of this strategy varies).

**Figure 3.**
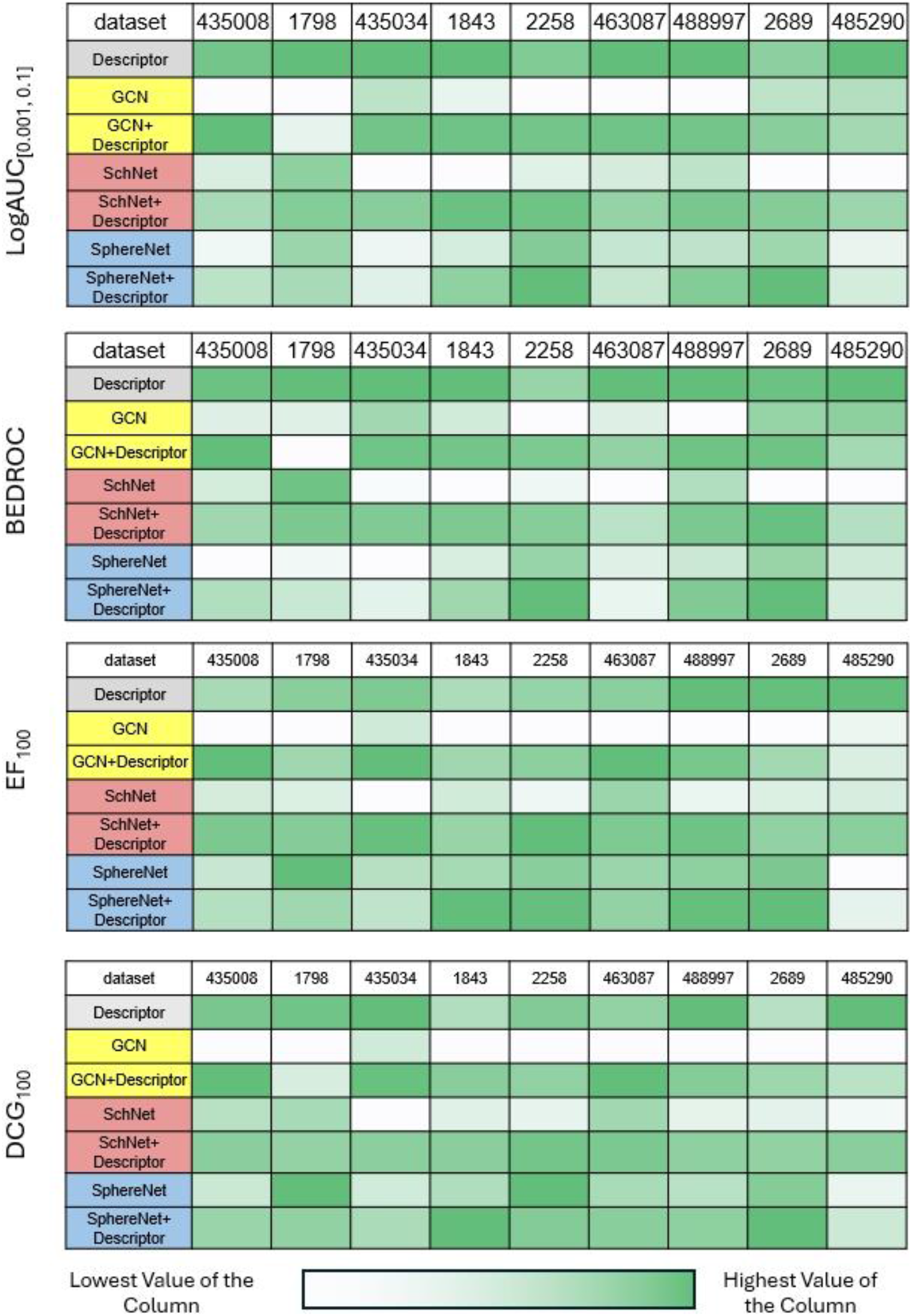
Scaffold Split: Performance of different models. The concatenation strategy still enhances the GNNs for most cases. Notably, descriptors perform better than many models across different metrics, especially salient in logAUC_[0,001,0,1]_ and BEDROC.

Most interestingly, we found that the descriptors alone outperform many GNNs. In some cases, it even outperforms the integrated-version GNNs. We hypothesize that this could result from the fact that deep learning-based methods are more easily overfit to the training data and therefore will perform worse than the expert-crafted ones when the data distribution is shifted. This finding prompts us to reconsider whether data-driven methods alone, despite their growing popularity, are the best approach for real-world drug discovery campaigns. Moreover, this also shows that even when coupled with descriptors, the performance of the integrated model may decrease and not always offer benefits. Finally, this finding emphasizes the need for developing better frameworks that integrate domain knowledge for improved predicted power under scaffold split scenarios.

## 3. Method

### 3.1. Datasets

We validate the effectiveness of the proposed strategy via nine well-curated high-throughput screening (HTS) datasets. To avoid issues with experimental artifacts and high false positive rates ^24^, for the validation of our strategy, we chose datasets carefully curated ^25^ from high throughput screens in the PubChem database ^26^. Only datasets with robust secondary validation of compounds were considered. Datasets details are shown in Table 1.

**Table 1.**
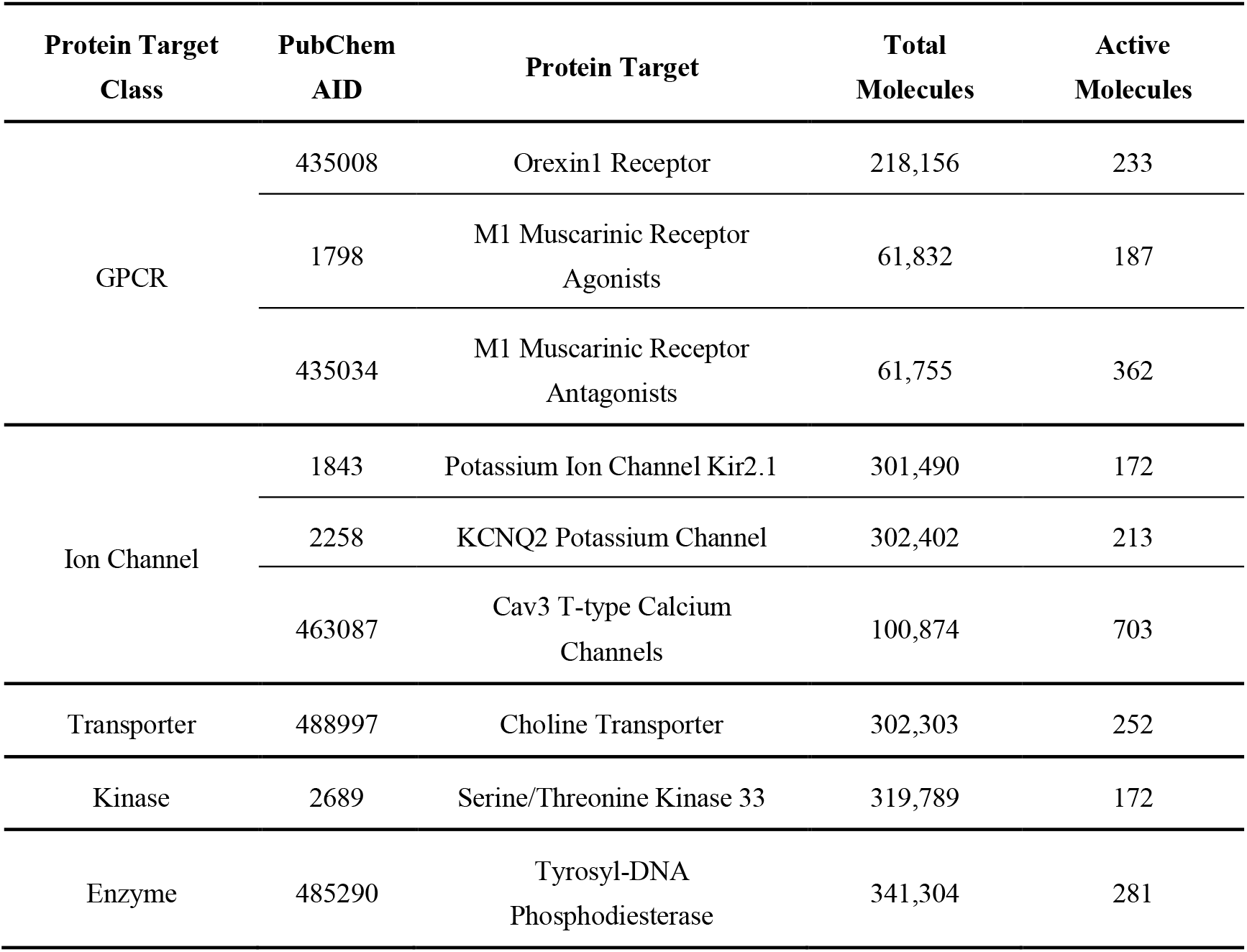
Dataset statistics.

| Protein Target Class | PubChem AID | Protein Target | Total Molecules | Active Molecules |
| --- | --- | --- | --- | --- |
| GPCR | 435008 | Orexin1 Receptor | 218,156 | 233 |
|  | 1798 | M1 Muscarinic Receptor Agonists | 61,832 | 187 |
|  | 435034 | M1 Muscarinic Receptor Antagonists | 61,755 | 362 |
| Ion Channel | 1843 | Potassium Ion Channel Kir2.1 | 301,490 | 172 |
|  | 2258 | KCNQ2 Potassium Channel | 302,402 | 213 |
|  | 463087 | Cav3 T-type Calcium Channels | 100,874 | 703 |
| Transporter | 488997 | Choline Transporter | 302,303 | 252 |
| Kinase | 2689 | Serine/Threonine Kinase 33 | 319,789 | 172 |
| Enzyme | 485290 | Tyrosyl-DNA Phosphodiesterase | 341,304 | 281 |

SMILES from the datasets were converted to SDF files using Open Babel ^27^. Standardized 3D coordinates are generated using Corina ^28^. Molecules are further filtered with atom type validity and duplicates with the BioChemical Library (BCL) ^21^.

Random split is used for the experiments, and each dataset is split into 80% for training and 20% for testing. Because preliminary results and previous literature ^19^ have shown that dropout can help avoid overfitting and the number of known active compounds is limited, we take the model from the last training epoch instead of the one from early stopping determined by validation performance. Multiple splits are used to prove the robustness of the proposed strategy.

### 3.2. Evaluation Metric

1. Logarithmic Receiver-Operating-Characteristic Area Under the Curve with the False Positive Rate in the range [0.001, 0.1] (**logAUC**_**[0.001,0.1]**_) Ranged logAUC ^29^ is used because only a small percentage of molecules predicted with high activity can be selected for experimental tests in consideration of cost in a real-world drug discovery campaign ^24^. This high decision cutoff corresponds to the left side of the Receiver-Operating-Characteristic (ROC) curve, i.e., those False Positive Rates (FPRs) with small values. Also, because the threshold cannot be predetermined, the area under the curve is used to consolidate all possible thresholds within a certain small FPR range. Finally, the logarithm is used to bias towards smaller FPRs. Following prior work ^19^, we choose to use logAUC_[0.001,0.1]_. A perfect classifier achieves a logAUC_[0.001,0.1]_ of 1, while a random classifier reaches a logAUC_[0.001,0.1]_ of around 0.0215, as shown below:

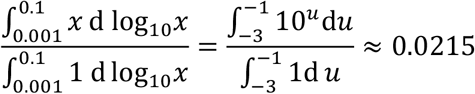
2. Boltzmann-enhanced discrimination of receiver operating characteristic (**BEDROC**) BEDROC ^25^ is a metric that evaluates the early recognition ability of a given model. It prioritizes the identification of active compounds early in the ranked list. BEDROC ranges from 0 to 1, where a score closer to 1 indicates better performance in recognizing active compounds early in the list.
3. Enrichment factor with cutoff 100 (**EF**_**100**_) Enrichment factor ^26^ is often used metric in virtual screening. It measures how well a screening method can increase the proportion of active compounds in a selection set, compared to a random selection set. Here we select the top 100 compounds as the selection set. And the EF_100_ can be defined as follows.

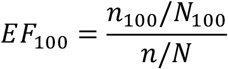 Where *n*_100_ is the number of true active compounds in the ranked top 100 predicted compounds given by the model, *N*_100_ is the number of compounds in the top 100 predicted compounds (i.e., 100), *n* is the number of active compounds in entire dataset, *N* is the number of compounds in the entire dataset. It is essentially a measure of the method’s ability to “enrich” the set of compounds for further testing. A random selection set receives a EF_100_ of 1. If no true active compounds are in the top 100 compounds, the EF_100_ becomes 0.
4. Discounted cumulative gain with cutoff 100 (**DCG**_**100**_) DCG ^27^ is a measure of ranking quality often used in web search. In a web search, it is obvious that a method is better when it positions highly relevant documents at the top of the search results. Virtual screening has a similar evaluation logic where we desire the active molecules to appear at the top of the selection set. To calculate DCG, a simpler version metric named cumulative gain (CG) ^27^ is introduced below. CG is the sum of the relevance value of a compound in the selection set. In our case, a true active compound receives a relevance value of 1, while a true inactive compound receives a relevance value of 0. So, the CG with cutoff 100 (CG_100_) equals the number of true active compounds in the top 100 compounds, i.e.,

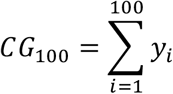

It can be observed that CG_100_ is unaffected by changes in the ordering of compounds. DCG hence aims to penalize a true active molecule appearing lower in the selection set by logarithmically reducing the relevance value proportional to the predicted rank of the compound, i.e.,

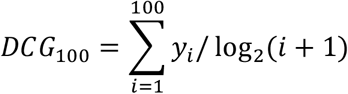

### 3.3. Baseline Models

We used three GNN models in our experiments: GCN ^20^, SchNet ^11^ and SphereNet ^13^. We used the BCL ^21^ to generate traditional QSAR descriptors. Following previous examples ^19, 30^, we use the optimal descriptors where 391-element molecular-level features are generated. We provide a brief introduction to each of the models and the BCL below.

GCN extends the concept of convolution from regular, grid-like data (such as images) to graphs, which have arbitrary structures. GCNs work by aggregating information from a node’s neighbors (potentially the node itself) to learn a representation of each node that captures both its features and local topology.

SchNet is a GNN designed for processing 3D molecules. The core design is continuous filters that are capable of handling unevenly spaced data, particularly, atoms. It also contains blocks that model interactions between atoms in a molecule.

SphereNet incorporates unique spherical message passing (SMP) for processing 3D molecules. The is encoded in a spherical coordinate system consisting of distance, angle and torsion. The SMP then uses the spherical coordinate system for the message passing process.

BCL is an application-based, open-source software package that integrates traditional small molecule cheminformatics tools with machine learning-based quantitative structure-activity/property relationship (QSAR/QSPR) modeling. It is designed to facilitate various cheminformatics tasks such as computing chemical properties, estimating druglikeness etc. It serves as a valuable resource for researchers in the computer-aided drug discovery field by providing a modular toolkit that supports the integration of cheminformatics and machine learning tools into their research workflows.

## 4. Future Work

In future work, we plan to expand our investigation by incorporating a broader array of GNN architectures and descriptor sets. This expansion will allow us to evaluate the generalizability and scalability of our integrative approach across a wider spectrum of computational models and chemical descriptor libraries.

We aim to explore advanced GNN models that may offer distinct advantages in capturing molecular features and interactions, potentially leading to improved predictive performance in virtual screening tasks. By comparing a diverse range of GNN architectures, we can better understand the nuances of how different models interact with various descriptor sets, and identify optimal combinations that maximize screening efficacy and accuracy.

Additionally, we intend to experiment with an expanded set of expert-crafted descriptors, including those that capture more intricate chemical and physical properties of molecules. This will enhance our ability to assess the impact of different types of descriptors on the performance of GNNs in virtual screening. By systematically evaluating the contribution of each descriptor type, we can refine our integration strategies to leverage the strengths of both GNNs and traditional chemical descriptors effectively.

Ultimately, our goal is to develop a comprehensive framework that can adapt to the evolving landscape of drug discovery, accommodating new advances in machine learning and cheminformatics.

## 5. Conclusion

Our study has rigorously evaluated the impact of integrating expert-crafted descriptors with GNNs and demonstrated that this integrative approach can significantly enhance the predictive power of virtual screening processes. Notably, the use of descriptors in conjunction with GNN architectures like GCN and SchNet has led to substantial improvements in identifying bioactive compounds.

In addition, The convergence in performance metrics across different GNN models, when supplemented with descriptors, suggests the potential for simpler GNN architectures to achieve results comparable to their more complex counterparts within this integrative framework. This finding underscores the viability of leveraging traditional knowledge and computational simplicity to advance the state-of-the-art in virtual screening.

Furthermore, our experiments with scaffold split scenarios revealed the robustness of descriptors, often outperforming combined GNN-descriptor models. This highlights the enduring value of expert knowledge in the face of evolving computational techniques and stresses the necessity for future models to effectively integrate this knowledge to enhance predictive power in realistic drug discovery settings.

In conclusion, our study serves as a compelling demonstration of how the synergistic integration of GNNs and expert-crafted descriptors can significantly advance the field of ligand-based virtual screening. As we move forward, it is imperative that we continue to explore and refine these integrative strategies, with the aim of developing more sophisticated and effective tools for drug discovery. The journey towards optimizing virtual screening methodologies is far from complete, but our work provides a significant step forward, offering a blueprint for future research in this dynamic and evolving field.

## Supporting information

protocol_capture

## Acknowledgments

Yunchao (Lance) Liu acknowledges that the Nvidia Academic Hardware Grant provides an A6000 GPU for speeding up the computation. Yunchao (Lance) Liu thanks Holy Gagnon for inspiring the discussion of the results.

## Appendix A. Descriptor Features

The descriptor sets used in this study are from ^19^. There are 391 elements of features in total. Each signed 2D autocorrelation (2DA ^31^) contains 32 bins. Each signed 3D autocorrelation (3DA ^31^) contains 60 bins. See the original paper ^19^ for individual feature naming details.

**Table 2.**
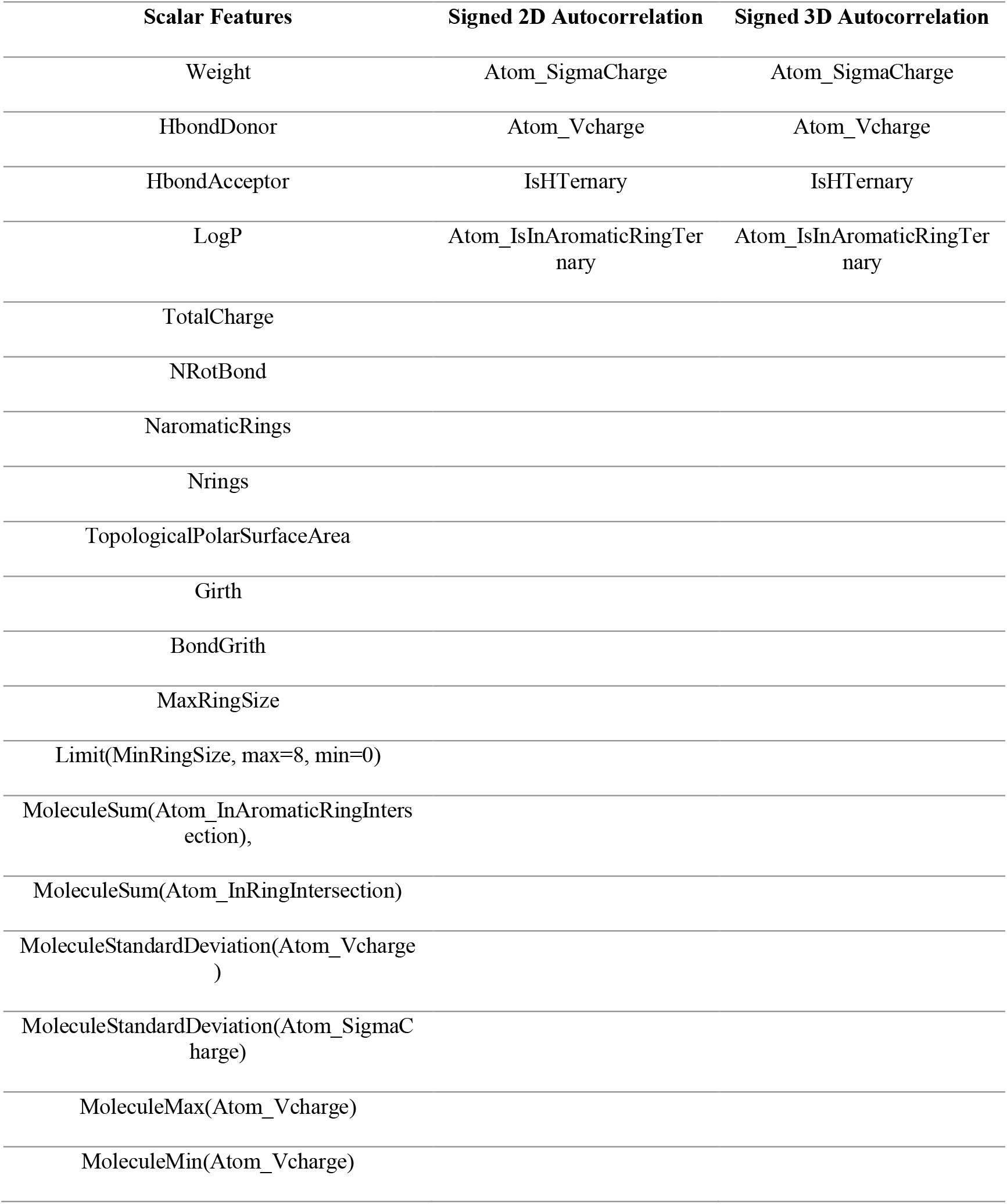
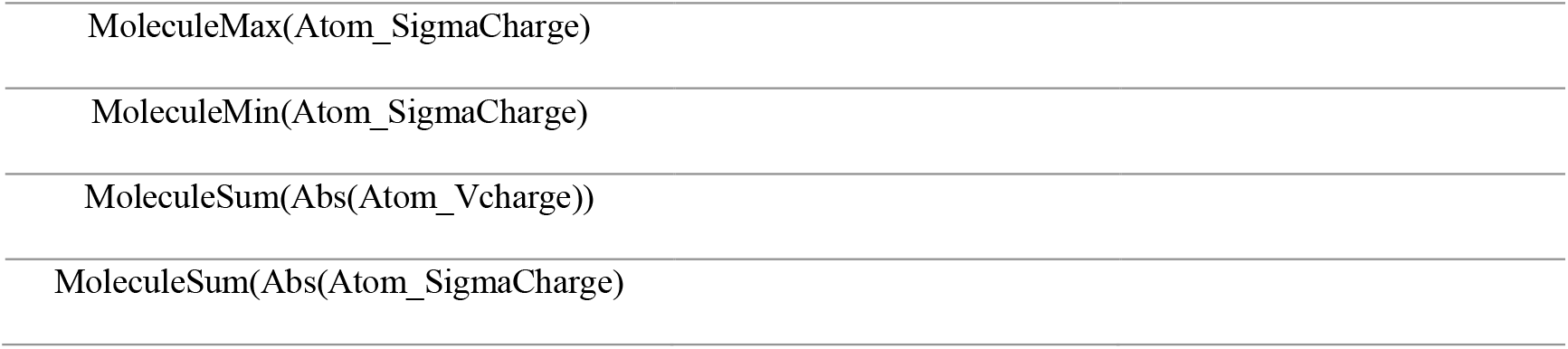
Features used in the descriptor set. Originally used in ^19^.

| Scalar Features | Signed 2D Autocorrelation | Signed 3D Autocorrelation |
| --- | --- | --- |
| Weight | Atom_SigmaCharge | Atom_SigmaCharge |
| HbondDonor | Atom_Vcharge | Atom_Vcharge |
| HbondAcceptor | IsHTernary | IsHTernary |
| LogP | Atom_IsInAromaticRingTernary | Atom_IsInAromaticRingTernary |
| TotalCharge |  |  |
| NRotBond |  |  |
| NaromaticRings |  |  |
| Nrings |  |  |
| TopologicalPolarSurfaceArea |  |  |
| Girth |  |  |
| BondGrith |  |  |
| MaxRingSize |  |  |
| Limit(MinRingSize, max=8, min=0) |  |  |
| MoleculeSum(Atom_InAromaticRingIntersection), |  |  |
| MoleculeSum(Atom_InRingIntersection) |  |  |
| MoleculeStandardDeviation(Atom_Vcharge) |  |  |
| MoleculeStandardDeviation(Atom_SigmaCharge) |  |  |
| MoleculeMax(Atom_Vcharge) |  |  |
| MoleculeMin(Atom_Vcharge) |  |  |

|  |
| --- |
| MoleculeMax(Atom_SigmaCharge) |
| MoleculeMin(Atom_SigmaCharge) |
| MoleculeSum(Abs(Atom_Vcharge)) |
| MoleculeSum(Abs(Atom_SigmaCharge)) |

