## Supplementary material for "Advancements in Ligand-Based Virtual Screening through the Synergistic Integration of Graph Neural Networks and Expert-Crafted Descriptors": protocol_capture

July 11, 2024

### 1 Prerequisites

#### 1.1 Datasets

High quality datasets from [1, 2] are used for the experiments. See original papers for details of datasets.

Datasets can be downloaded from `https://figshare.com/articles/dataset/Well-curated_QSAR_datasets_for_diverse_protein_targets/20539893`

Decompress the downloaded file, to get .sdf files.

#### 1.2 Creating BCL Features

This section will process the datasets described in the previous section into BCL features. The output of this section is a new folder named BCL-feats and BCL features are stored in (dataset-BCL-feat.csv) in that folder.

BCL can be downloaded at `https://github.com/BCLCommons/bcl/`. See `https://www.frontiersin.org/articles/10.3389/fphar.2022.833099/full` for an introduction of BCL. The following scripts assume the *bcl.exe* command points to a working BCL program.

After downloading the scripts (.py) and configuration files (.obejct) from `https://github.com/meilerlab/gnn-descriptor/BCL`, place them according to Fig. 1.

When at the BCL folder, execute the following commands step by step to generate BCL features:

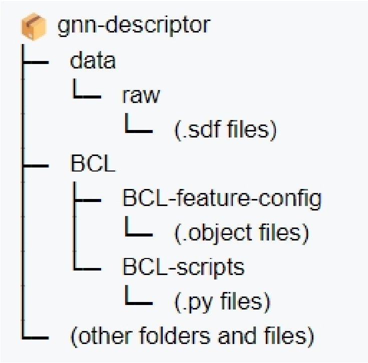


Figure 1: In order to run the codes, the files should be named and placed according to this diagram. The datasets (.sdf files) should be placed under *raw* folder. The BCL configuration files (.object) should be placed under *BCL-feature-config* folder. The BCL-scripts should be placed under *BCL-scripts* folder. All the other files and folders from the github repository are placed as they are in the gnn-descriptor folder.

```
python BCL-scripts/1_combine_sdf.py # This combines active and
    inactive SDF files into one SDF file.
```

```
python BCL-scripts/2_add_id_to_sdf.py # This will add IDs to the
    header of each molecule in the SDF. So it is easy to identify
    which molecules are filtered out in the later steps.
```

```
python BCL-scripts/3_filter.py # Add hydrogen, neutralize molecules
     and filter out molecules that do not have simple atoms (C, O,
    N, S, P, F, Cl, Br, I)
```

```
python BCL-scripts/4_data_preparation.py  # This creates the BCL
    features and stores that in the BCL-feats folder.
```

```
python BCL-scripts/5_count_unmatched.py # This counts how many
    molecules are filtered out.
```

```
python BCL-scripts/6_clean_intermediate.py  # This cleans up the
    generated intermediate files to save hardware space. ‘
```

### 2 Running the Codes

#### 2.1 General Instructions

Install the following python library:

- pytorch 2.0.1
- torch-geometric 2.3.0
- pytorch-scatter 2.1.2
- rdkit 2023.9.1
- tqdm
- pandas 2.1.2
- numpy 1.26.1

All necessary scripts can be downloaded at `https://github.com/meilerlab/gnn-descriptor`

To run the codes, the first thing is to set the running parameters as a .cfg file in the config folder. Below, an example of a GCN configuration file is shown as *GCN.cfg*

```
[GENERAL]
seed = 1
num_workers = 12

[DATA]
dataset_name = 1798
root = data
split_scheme = random1_IAratio100_cv0

[TRAIN]
num_epochs = 50
batch_size = 32
warmup_iterations = 2000
peak_lr = 1.4e-4
end_lr = 1e-9


[MODEL]
model_type = gcn # all model names should be lowercased
in_channels = 28
hidden_channels = 32
num_layers = 4
with_bcl = False
bcl_dim = 391
```

The experiments in the main texts used split_scheme = ['random1_IAratio100_cv0', 'random1_IAratio100_cv1', 'random1_IAratio100_cv2'] and mode_type = ['mlp', 'gcn', 'schnet', 'spherenet']

Example command:

```
python main --config config/GCN.cfg --no_train_eval --test
```

The *GCN.cfg* should be replaced with a configuration file specified by the user.

–*config* is a required flag and it specifies the configuration file.

–*no_train_eval* is an optional flag. When the flag is added, the training evaluation is skipped.

–*test* is an optional flag. When the flag is added, the training is skipped and a saved model (in *saved_model* folder) is directly evaluated at the testing set.

#### 2.2 Use Cases

##### Use User-Specified Data

Place the .sdf files in the *raw* folder in Fig. 1. There should also be a file specifying how the data is split into train and test. The split file is put into *split* folder. The split file is a .pt file that contains a python dictionary. The dictionary has two keys, 'train' and 'test'. The value for either key is a list of index numbers of the data.

A example split.pt file: {'train':[1,3,8,...2943], 'test':[2,4,5,6,7, 2944]}

##### Train a Model

To train the model scratch, first set the model.cfg configuration file and placing that in the *config* folder as shown in Sec. 2.1, you can train the model with

```
python main --config config/model.cfg
```

*If you do not want to see the evaluation metrics during metrics, –*no_train_eval* flag can be used to speed up the training.

##### Test a Pre-trained Model

First, place the pre-trained model in the *saved_model* folder. Then the model can be run with the test model with the following command:

```
python main --config config/model.cfg --test
```

The –*test* flag helps to skip the training process and directly evaluate the model
